# Decoding molecular mechanisms for loss-of-function variants in the human proteome

**DOI:** 10.1101/2024.05.21.595203

**Authors:** Matteo Cagiada, Nicolas Jonsson, Kresten Lindorff-Larsen

## Abstract

Proteins play a critical role in cellular function by interacting with other biomolecules; missense variants that cause loss of protein function can lead to a broad spectrum of genetic disorders. While much progress has been made on predicting which missense variants may cause disease, our ability to predict the underlying molecular mechanisms remain limited. One common mechanism is that missense variants cause protein destabilization resulting in decreased protein abundance and loss of function, while other variants directly disrupt key interactions with other molecules. We have here leveraged machine-learning models for protein sequence and structure to disentangle effects on protein function and abundance, and applied our resulting model to all missense variants in the human proteome. We find that approximately half of all missense variants that lead to loss of function and disease do so because they disrupt protein stability. We predicted functionally important positions in all human proteins and found that they cluster on protein structures and are often found on the protein surface. Our work provides a resource for interpreting both predicted and experimental variant effects across the human proteome, and a mechanistic starting point for developing therapies towards genetic diseases.

## Introduction

Proteins are responsible for the majority of cellular functions, and perform most of these functions by interacting with other molecules. These molecular interactions occur at sites or regions whose identification is critical to understanding human biology and disease (***Yue et al., 2005a***; ***Sahni et al., 2015***; ***Starita et al., 2017***). Experimentally, such functionally important positions are often identified by examining how amino acid changes affect protein function, stability, and interactions (***Starita et al., 2017***).

In addition to their importance in mutational analyses, missense variants—in which a single amino acid residue is replaced by another—are estimated to account for between one-third and one-half of all pathogenic variants currently catalogued in clinical databases (***Taipale, 2019***). While a large fraction of missense variants in a given protein are either well tolerated or have a modest effect on protein functionality (***Cagiada et al., 2021***), the subset that have a strongly deleterious effect can cause loss of function by a variety of mechanisms; they may for example directly affect protein function or stability, subtly disrupt function by altering protein folding pathways, increase degradation rates, or affect the ability of proteins to interact with their environment (***Sánchez et al., 2006***; ***Gray et al., 2017***; ***Schaafsma and Vihinen, 2017***). Identifying deleterious variants and understanding their molecular mechanism underlying perturbed function is therefore important for understanding the role of each residue in the protein (***Chiasson et al., 2020***; ***Faure et al., 2021***; ***Cagiada et al., 2021***), the role of the protein itself (***Deák and Cook, 2022***), and for assessing potential pathogenicity (***Quinodoz et al., 2022***; ***Lin et al., 2023***). In the last decade, large-scale mutagenesis experiments, such as multiplexed assays of variant effects (MAVEs), have facilitated the analysis of thousands or even millions of missense mutations within a single experiment (***Gasperini et al., 2016***; ***Weile and Roth, 2018***), providing information on how individual residue substitutions may affect cellular abundance, stability, and function of the protein. Such experiments have provided a wealth of information about which substitutions may cause loss of protein function, and have served as critical data to benchmark prediction methods. Nevertheless, because most proteins need to fold in order to function, it is difficult to disentangle variants that directly perturb function from those that do so indirectly by affecting the stability and abundance of the protein structure (***Echave and Wilke, 2017***; ***Li and Lehner, 2020***).

Experimentally, it is possible to differentiate between molecular effects by examining multiple properties such as protein abundance and function. Moreover, MAVEs make it possible to scale these analyses to entire proteins. Inspired by these ideas, we recently developed a computational approach that combines analyses of protein structure and evolution to find functionally relevant residues, and for differentiating between variants that cause loss of function through either reduced abundance or by directly affecting protein features such as intermolecular interactions (***Cagiada et al., 2021***, ***2023***). Briefly described, we performed computational saturation mutagenesis using (i) a method to assess variant effects on protein stability and (ii) a method to assess the impact of a variant in terms of the evolutionary record encoded using a multiple sequence alignment. In this way, we used the evolution-based models to predict loss of protein function, and the stability predictions to find those loss-of-function variants that are caused by perturbed protein stability and function. We also showed that residues that are conserved in evolution, but not due to a role for protein stability, are highly enriched in verified functionally relevant residues, and validated the results using prospective experiments in HPRT1 (***Cagiada et al., 2023***). Our approach thereby provided proof of principle for the ability to simultaneously predict disease variants and the molecular mechanisms underlying variants effects, and to discover functionally important positions in proteins (***Cagiada et al., 2021***, ***2023***).

While this approach was accurate it relied on relatively slow computational methods for assessing variant effects, including the construction of high-quality multiple sequence alignments, limiting proteome-wide applications. Further, our approach was based on training a machine-learning model on a small set of proteins that have been studied experimentally by multi-modal MAVEs. To address these issues and enable accurate proteome-wide analyses we here present an approach that leverages accurate and fast pre-trained machine-learning models. In recent years, protein language models have been shown to enable accurate predictions of variant effects that approach the accuracy of alignment-based methods (***Meier et al., 2021***; ***Outeiral and Deane, 2022***; ***Brandes et al., 2023***) and for assessing the impact of variants on protein stability (***Notin et al., 2023***). Specifically, we here combine the ESM-1b model (***Rives et al., 2021***) for protein ‘conservation’ with the ESM-IF (‘Inverse Folding’) model (***Hsu et al., 2022***) to assess the impact of variants on protein stability via a model that we call FunC-ESMs (Functional Characterisation via Evolutionary Scale Models). We validate FunC-ESMs across a range of scenarios and apply it to characterise all possible missense variants in the human proteome including seventy thousand clinically annotated variants distributed across more than ten thousand proteins.

## Results and Discussion

### FunC-ESMs: a functional characterisation model based on ESM representations

We developed FunC-ESMs to provide a rapid and accurate approach to (i) predict which variants may cause loss of function, (ii) find those variants that cause loss of function via decreased protein stability, and (iii) use these predictions to discover functionally relevant residues. FunC-ESMs builds on our previously described ideas (***Cagiada et al., 2021***, ***2023***) and extends these by (i) avoiding the need for accurate sequence alignments, (ii) avoiding possible biases from training a machine learning on a small dataset, and (iii) performs predictions at substantially improved speeds. In previous work we used either Rosetta (***Park et al., 2016***) or RaSP (***Blaabjerg et al., 2023***) to assess stability effects; here we instead use ESM-IF, which combines a graph neural network and a language model to represent proteins (***Hsu et al., 2022***). It has been shown (***Notin et al., 2023***) that ESM-IF representations can be used to predict the effects of variants on protein stability with accuracy that is comparable to, or sometimes surpasses, that of traditional energy-based predictors. Moreover, these predictions can be obtained in just a few seconds, even for large proteins. As input for ESM-IF, we used the backbone of structures generated with AlphaFold2 (AF2; ***Jumper et al.*** (***2021***)), which provides a complete residue coverage and with a quality that is sufficient for a wide range of structural bioinformatics tasks (***Akdel et al., 2022***). To assess sequence conservation we previously applied GEMME (***Laine et al., 2019***), which uses sequence alignments to score variant effects; here we instead use the protein language model ESM-1b (***Rives et al., 2021***) which provides comparable accuracy for variant effects (***Meier et al., 2021***; ***Brandes et al., 2023***) at substantially improved speeds. As described in more detail below, FunC-ESMs combines information from ESM1b and ESM-IF to predict and assign mechanisms to variant effects (Fig. 1).

**Figure 1.**
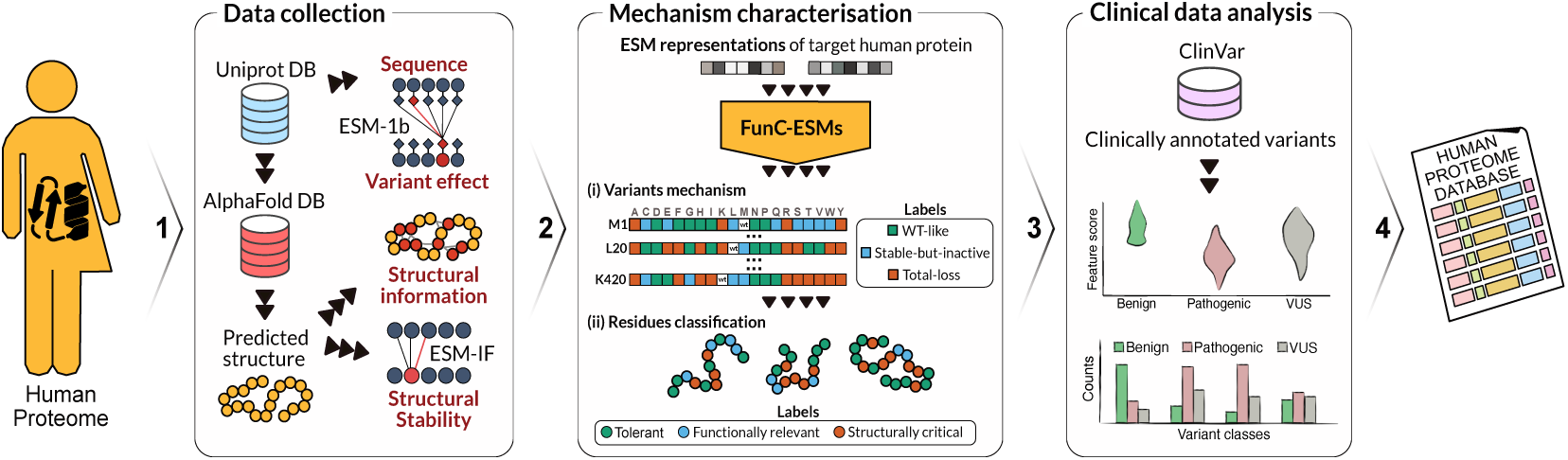
Graphical summary of our functional characterisation of the human proteome. (1) For all human proteins, we collected the corresponding sequence and predicted structure. We used these as input to ESM-1b and ESM-IF to generate variant effect scores probing conservation and stability. (2) We combined these data as input to FunC-ESMs which assigns an output label to each variant. We used these labels to further assign a functional annotation for all residues. (3) We collected clinical annotations from ClinVar. We analysed these in light of the different ESM scores and used FunC-ESMs to provide insight into the mechanism of loss of function. (4) The resulting information is made available in an online database at the Electronic Research Data Archive at University of Copenhagen (KU/UCPH) (ERDA) available via https://sid.erda.dk/cgi-sid/ls.py?share_id=DUWFpyjZp0. The results can also be accessed with a colab notebook available at https://github.com/KULL-Centre/_2024_cagiada-jonsson-func.

Because ESM-1b and ESM-IF directly provide accurate (so-called ‘zero-shot’) predictions, we used their scores directly as input to a threshold-based method, thus circumventing the need for training a supervised model on a relatively small set of proteins. FunC-ESMs first assigns a ‘deleterious’ label to a variant based on its ESM-1b score and then, for those variants labelled deleterious, assesses the effect on structure and function based on the ESM-IF score. Thus, each variant is classified as either ‘WT-like’ (non-deleterious), or as ‘total-loss’ (important for stability and therefore for function) or ‘stable-but-inactive’ (important for function but not stability). We optimized the threshold used for ESM-1b by using data from 32 MAVEs in ProteinGym (***Notin et al., 2022***) and obtained an optimal threshold for ESM-1b to separate WT-like and deleterious variants (Fig. S1A,C). For ESM-IF, we used nearly two hundred thousand ΔΔ*G* values (***Tsuboyama et al., 2023***) to select a threshold for the ESM-IF score to separate stable and unstable variants (Fig. S1B,D).

We then validated FunC-ESMs using both the data that we previously used to train a model (***Cagiada et al., 2023***) as well as labels obtained from experimental measurements on the SH3 domain of Growth Factor GRB2 (***Faure et al., 2021***) and human glucokinase (***Gersing et al., 2023a***,b). For all five proteins we find a high prediction accuracy comparable to our previous model (Figs. S2 and S3), thus demonstrating the accuracy of FunC-ESMs across a range of molecular functions that include both enzymes and proteinprotein interactions. These results confirm the utility of using the computationally efficient and accurate ESM-1b and ESM-IF models to predict functionally relevant residues and classify variant effects. In addition to running FunC-ESMs on the entire human proteome (see below), we also make the code available, and provide easy access to running the model via Google Colaboratory (see Methods).

### A functional characterization of variants and residues in the human proteome

Having assessed the accuracy of the FunC-ESMs to identify the mechanism behind loss-of-function variants, we proceeded to predict variant effects and assign residue labels for the proteins in the human genome and the mechanism behind disease variants (Fig. 1). We generated ESM-1b and ESM-IF scores, using as reference one protein sequence per human gene from the UniProt human reference proteome. We collected the predicted structures from the AlphaFold2 protein database (***Jumper et al., 2021***; ***Varadi et al., 2022a***) and used these as input for ESM-IF. Similarly, we used the sequence for all human proteins as input to ESM-1b to assess variant effects. To analyse and interpret the output from FunC-ESMs, we also collected predictions of protein disorder (***Hanson et al., 2017***; ***Alderson et al., 2023***), and calculated descriptors of the biochemical environment and residue exposure. In total, we collected data for 20,144 human proteins and to make predictions with FunC-ESMs on 195,780,218 variants, distributed over 10,304,222 residues.

The model predictions were used to designate each individual variant as either wildtype-like (WT-like), total-loss (TL), or stable-but-inactive (SBI). We found that 60% of the predicted substitutions were classified as WT-like, 19% as total-loss and 21% as stable-but-inactive (Fig. S4A). At the residue level, we also assigned a label based on the role of the residue in the protein: we designated residues where most variants were WT-like as ‘tolerant’; and refer to positions critical for maintaining protein stability (with a dominance of total-loss variants) as ‘structurally critical’. Residues with many variants predicted to be stable-but-inactive were labelled as ‘functionally relevant’; residues without any dominant effect were designated as ‘mixed’. Our results show that 57% of all residues were labelled as tolerant, 17% as structurally critical, 19% as functionally relevant, and 7% as mixed (Fig. S4D).

The partitioning between different classes of residues was similar to our previous analyses using both experiments and computation on two proteins (***Cagiada et al., 2021***). As these previous analyses were performed on globular and folded regions, we separated the residues based on whether they were in a folded or intrinsically disorder region (IDR). Of the 10,304,222 residues that we analysed, we could label 10,093,187 as either folded (6,755,767; 67%) or disordered (3,337,420; 33%). As expected, the results differed substantially between residues in folded and disordered regions (Fig. S4). In the folded regions, there was a roughly equal amount of WT-like substitutions (48%) and deleterious variants (52%) (Fig. S4B,E). Among these deleterious variants, 52% were predicted to affect folding stability and 48% to cause loss of function via other mechanisms. In contrast, we found that the majority of the variants (85%) were predicted to be WT-like in the IDRs (Fig. S4C,F) in line with the known increased tolerance of IDRs to amino acid variation. Of the 16% of variants that are predicted to cause of loss of function, FunC-ESMs assigns most (94%) as affecting function, though we note that AF2 structures do not generally represent IDRs accurately (***Ruff and Pappu, 2021***; ***Tesei et al., 2024***). Nevertheless, disordered regions for which AF2 predicts local structuring may represent residues that are conditionally folded (***Alderson et al., 2023***) and are known to have increased fractions of pathogenic missense variants (***Alderson et al., 2023***; ***Tesei et al., 2024***).

We then focused on further characterising the residues in the folded regions of the human proteome. Our initial investigation examined whether, in proteins where more than half of the residues are folded, the fraction of residues with a deleterious effect on protein function correlates with protein size. We found that the fraction of residues that are predicted to be functionally relevant increases with protein size from 3% for small proteins (less than 100 residues), to 8% for proteins with a size corresponding to one to two domains, to 10% and 15% for larger presumably multi-domain proteins (Fig 2A). We found a similar trend for structurally critical residues (Fig. 2B). Given the differences among the 20 common amino acids we asked whether they contribute differently to different classes of residues (Fig. S5A,B). As expected, approximately 56% of the structurally critical residues in folded regions are aliphatic or aromatic, and these residues are depleted of charged amino acids. In contrast, polar and charged residues are enriched in functionally relevant residues in folded regions compared to structurally critical positions, consistent with the fact that polar residues are located on protein surfaces and are important for protein-protein interactions and enzyme catalysis. For cysteine residues, we also investigated their potential to participate in the formation of a disulfide bridge. Of the nearly 200,000 cysteine residues that we analysed, almost 50,000 were predicted to form disulphide bridges and of these, 80% were classified as structurally critical; in contrast only 28% of the cysteines not involved in disulphide bridges were predicted to be structurally critical.

**Figure 2.**
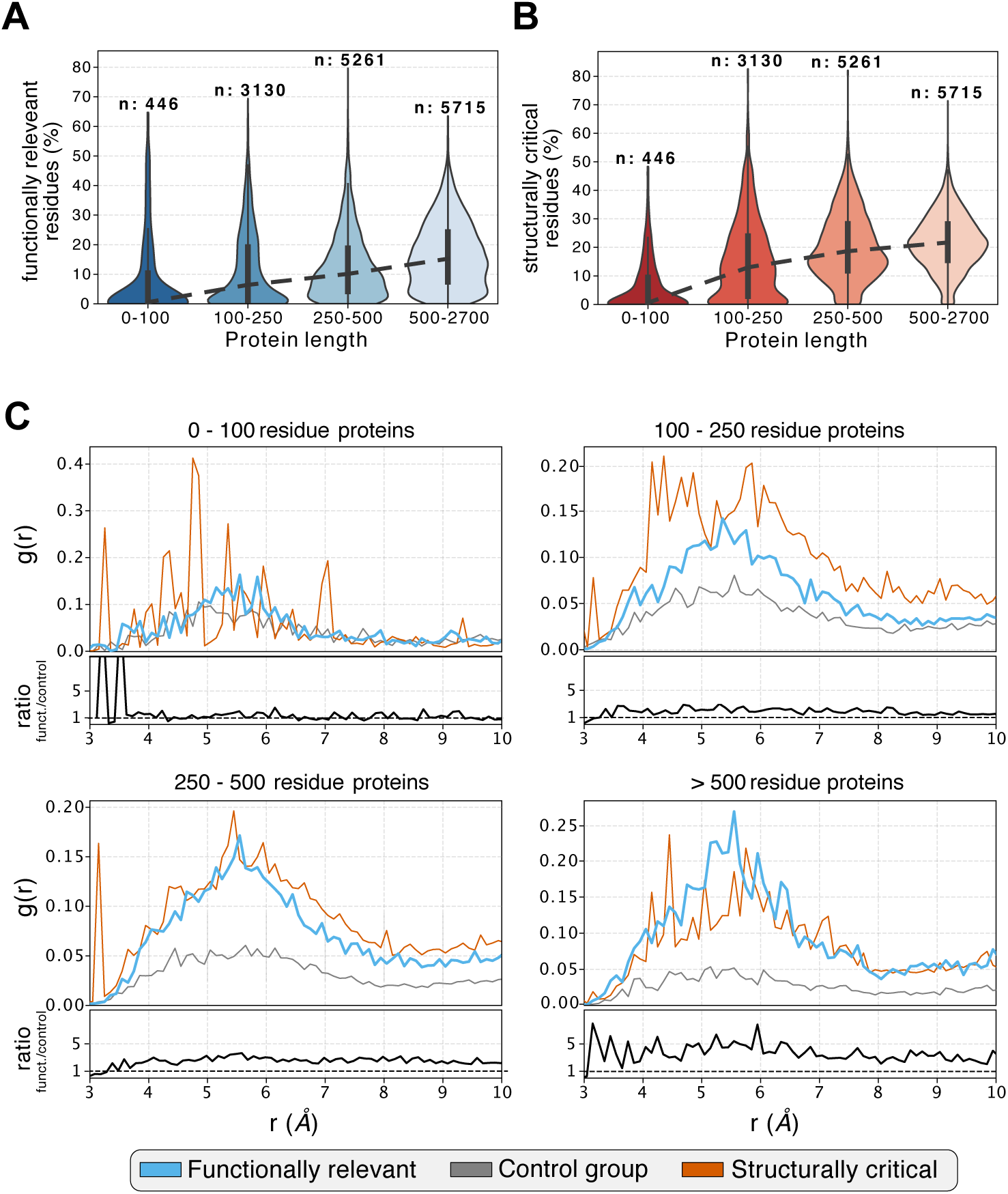
Analysis of functionally relevant and structurally critical residues in the human proteome. We compared the percentage of (A) functionally relevant and (B) structurally critical residues for proteins of different sizes. In each plot, the white dot in the centre of the distribution represents the median, the thick black bar represents the interquartile range and the thin black line represents the range of the data. The number of proteins in each subset is shown at the top of each violin plot. (C) Mean radial distribution functions (*g*(*r*)) for all analysed human proteins clustered by chain length. The blue lines represent *g*(*r*) for functionally relevant residues, the red line for the structurally critical residues, and the grey line for a control group. Below each plot we show in black the ratio of *g*(*r*) for the functionally relevant residues and the control group; values greater than one show that functionally relevant residues are more strongly clustered than the control group.

We then investigated whether certain types of amino acid substitutions generally had a greater impact on function or structure compared to others. (Fig. S5B,C). As expected, we found that substitutions from a hydrophobic to a polar or charged amino acid affect the stability of the structure more than substitutions to another hydrophobic amino acid. In line with previous observations (***Gray et al., 2017***; ***Dunham and Beltrao, 2021***; ***Høie et al., 2022***), we also found that mutations to proline are highly likely to disrupt the protein fold, regardless of the starting amino acid type. On the other hand, we did not observe strong signals for types of substitutions that cause loss of function without loss of stability.

Next, we examined whether the types of variant effects are distributed differently in residues that are either exposed or buried in the protein structure. Using our definition of burial (see Methods), we found that 38% of residues in the folded regions are buried in the protein structure, and that approximately two thirds of these positions fall in the either the ‘structurally critical’ or ‘functionally relevant’ class in which most substitutions have a deleterious effect (Fig. S6A). As expected, loss of function due to loss of stability is the dominant mechanism for loss of function at buried positions (Fig. S6A). Indeed, 80% of the structurally critical residues are located in the buried regions of the protein, while functionally relevant residues are mostly located in exposed regions (76%, Fig. S6B). We calculated the ‘half-sphere exposure’ (HSE; ***Hamelryck (2005)***) to analyse the surroundings of the different amino acids with ‘up’ referring to contacts in the direction of the side chain and ‘down’ in the direction of the backbone. This analysis showed that side chains at positions critical to folding stability typically reside in a more crowded protein environment than those at functionally relevant or tolerant positions, with more than twice the average HSE values of functionally relevant positions. In contrast, at the backbone level, the exposures are comparable between the different residue classes (Fig. S6C).

We then hypothesised that functionally relevant residues, owing to their critical and synergistic roles, would be spatially clustered. This hypothesis aligns with previous studies, which have demonstrated a spatial clustering for evolutionarily conserved residues (***Jack et al., 2016***; ***Mayorov et al., 2019***). To analyse co-localization of functionally relevant residues in the human proteome, we evaluated the radial distribution function (*g*(*r*)) for both functionally relevant and structurally critical residues in each folded region for proteins with more than half folded residues. Additionally, as a control, we calculated *g*(*r*) for a control group formed by randomly sampling an equivalent number of positions from the combined set of wild-type and functionally relevant residues. Briefly, *g*(*r*) quantifies the likelihood of finding other residues of the same type at a distance *r*. Therefore, peaks of *g*(*r*) at short distances suggest that residues of the same class are likely to be found in the vicinity of each other. We summarise the results by calculating the mean of all *g*(*r*) distributions for each class and comparing the resulting curves (Fig. 2C). While the results are noisy for the smallest proteins (less than 100 residues) the results for larger proteins show clearly that both structurally critical and functionally relevant residues are clustered in the protein structure. The clustering of structurally critical residues can be explained by their location in the hydrophobic and buried core (Fig. S6), whereas functionally relevant residues can be explained by their tendency to co-localise in clusters, such as in binding- or active sites. Indeed, we found that the radial distribution function for functionally relevant residues is consistently higher (showing greater clustering) than the control group.

### Investigating the mechanistic basis of disease variants in the human proteome

Having made predictions of the mechanistic consequences for all missense mutations in the human proteome, we next investigated the occurrence and characteristics of the mechanisms associated with the onset of disease for clinically annotated variants. To this end, we collected missense variants from ClinVar (***Landrum et al., 2014***) that were annotated as either (likely) ‘benign’ or ‘pathogenic’ (see Methods). In total, we extracted 102,056 clinically annotated human variants (69,909 benign and 32,147 pathogenic), distributed across 12,114 proteins. Prior to characterising the effects of these variants, we assessed the accuracy of the input features of FunC-ESMs (the ESM representations) as predictors of clinical pathogenicity for our proteomic dataset. This analysis aimed to reaffirm the already established accuracy of the machine-learning model as predictors of clinical pathogenicity (***Brandes et al., 2023***), by re-evaluating on a slightly different dataset. As expected, we find that the sequence-based model (ESM-1b) shows higher accuracy when predicting pathogenicity compared to the structure-based model (ESM-IF), with an area under the receiver operating characteristic curve (AUC) of 0.92 versus 0.83 (Fig. 3A). Thus, FunC-ESMs, which uses ESM-1b to divide variants into WT-like vs. deleterious, achieves close-to state-of-the-art accuracy in assessing pathogenicity (***Brandes et al., 2023***). This observation provides the starting point for our analysis of the molecular mechanisms for loss-of-function variants; specifically, we used FunC-ESMs to separate clinically annotated variants into the three classes (WT-like, stable-but-inactive, and total-loss) and compared these to their clinical status (benign or pathogenic) (Fig. 3B).

**Figure 3.**
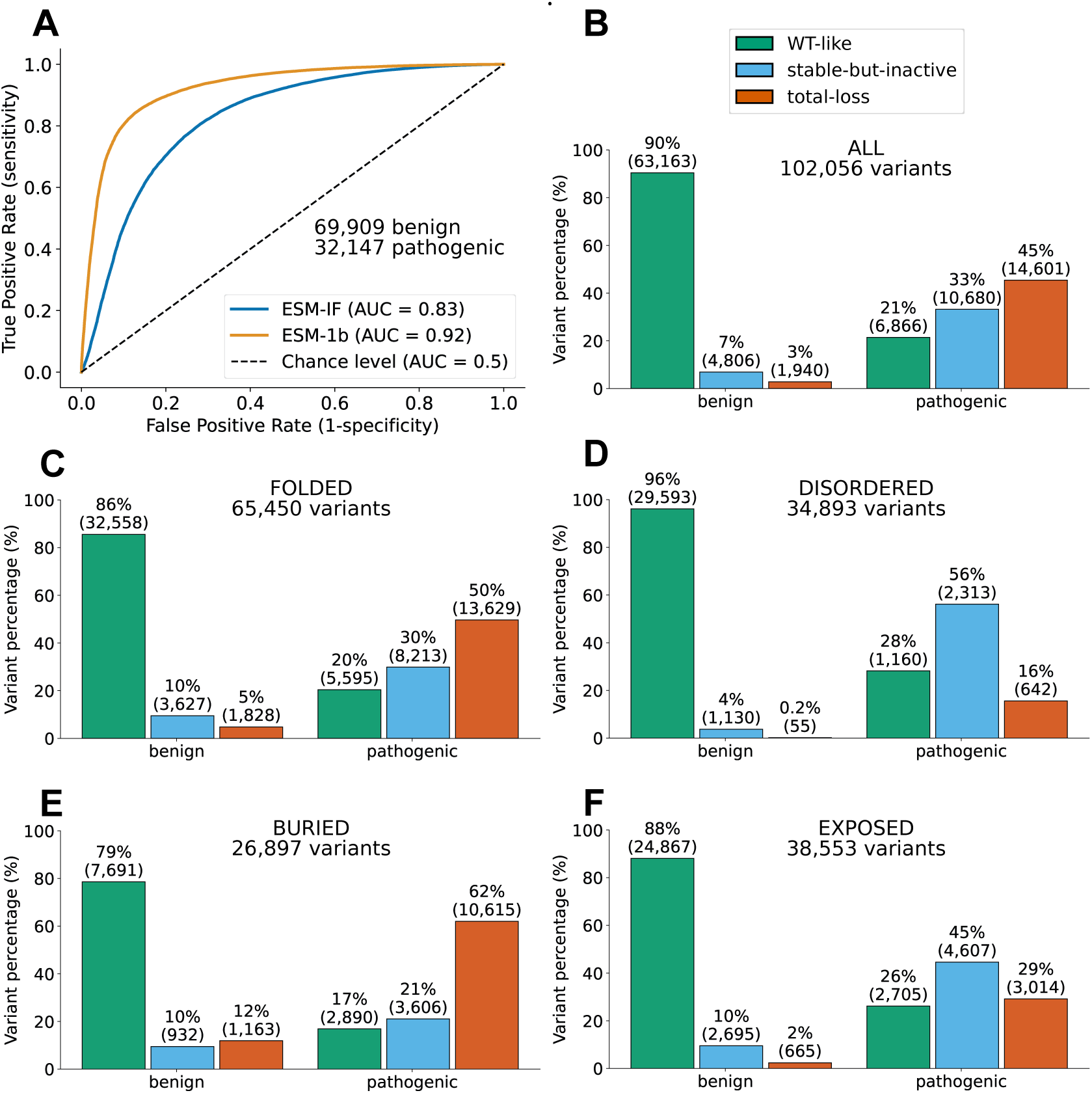
Molecular mechanisms of pathogenic variants. (A) Assessment of the accuracy of the two ESM models as predictors of clinical pathogenicity using receiver operating characteristic analysis. We used the ESM-1b and ESM-IF models to separate the clinically annotated ‘benign’ and ‘pathogenic’ missense variants and calculated the area under the curve (AUC) for each model. (B) Classification of benign and pathogenic missense variants using FunC-ESMs: ‘WT-like’ (green), ‘stable-but-inactive’ (blue), and ‘total-loss’ (red). We further divide these variants into subgroups depending on whether they are located in (C) folded or (D) intrinsically disordered regions, and whether they are (E) buried or (F) exposed. The total number of variants in each subset is shown at the top of each subplot for the different conditions. All bars are annotated with both the absolute counts and percentages for each category.

As expected from the accuracy of ESM-1b, we found that FunC-ESMs classifies most (90%) of the benign variants as WT-like while 78% of the pathogenic variants fall in either the stable-but-inactive or total-loss classes (Fig. 3B). While a small number of benign variants were incorrectly categorized as stable-but-inactive and as total-loss (7% and 3%, respectively), the overall division of variants into ‘WT-like’, ‘stable-but-inactive’, and ‘totalloss’ underlines the capacity of the model to differentiate the impacts of benign versus pathogenic variants. For pathogenic variants, our predictions classified about 78% of annotated variants as deleterious to protein function. Here we found that a large number of variants affect protein stability (45% of the total, or 58% of the correctly predicted pathogenic variants) and a smaller fraction that only affect function by other mechanisms (33% of the total or 42% of the correctly predicted variants). These results show that, across the human proteome, more than half of pathogenic missense variants cause loss of function via loss of protein stability, with the remaining directly affecting other aspects of protein function.

As discussed above, there are substantial differences in the patterns we observe for variants that are located in folded or disordered regions, and whether they are buried or exposed in the structure. Therefore, we also analysed the clinical variants in this context (Fig. 3C–F). In folded regions (Fig. 3C), a substantial majority (80%) of the pathogenic variants were classified as deleterious by FunC-ESMs, with about a third (30% of the total or 38% of the correctly predicted) being predicted to cause loss of function by directly impairing protein function, and the test being classified as total-loss (62% of the correctly predicted). This trend is in line with our expectations given the structural constraints in these regions. The results for intrinsically disordered regions (IDRs) present a contrasting picture (Fig. 3D), where we find that most pathogenic variants are predicted to cause loss of function by directly impairing protein function. We also note that the number of misclassified variants (false negative) is higher in IDRs (28% vs. 20%). This highlights the challenges in modelling variant effects in flexible regions, which are often poorly conserved by traditional measures (***Lindorff-Larsen and Kragelund, 2021***), and possibly that our cut-offs for the ESM-1b scores reflect MAVEs on mostly folded proteins. Given the conformational heterogeneity of IDRs and the notion that AF2 does not describe most conformational properties of IDRs, it is expected that loss of thermodynamic folding stability plays only a small role in IDRs. Indeed, only 16% of the loss-of-function variants are predicted to cause a loss of stability, a figure comparable to estimates suggesting that 15% of disordered residues may fold upon binding to their targets (***Alderson et al., 2023***), and in line with the finding that pathogenic variants are enriched in areas with high pLDDT scores within disordered regions (***Alderson et al., 2023***; ***Tesei et al., 2024***).

Finally, we explored the differences in variant effects between buried and exposed residues, focusing exclusively on those located within folded regions (Fig. 3E, F). The results are in line with the general effects across the proteome and expectations based the structure and chemistry of proteins. For pathogenic variants located at buried positions, 62% of those predicted as loss-of-function variants we predict to be caused by decreased protein stability. In contrast, among pathogenic variants at exposed positions, only 29% are predicted to result in loss of function due to reduced stability. The remaining 71% of these exposed pathogenic variants likely cause a direct loss of function by perturbing functional residues and intermolecular interactions. The increased prevalence of disease-associated variants in buried regions and a decreased prevalence in exposed regions highlight the differential impacts of residue location within protein structures on disease pathogenicity, and are in line with previous analyses (***Savojardo et al., 2021***). Overall, the ability of FunC-ESMs to discern the complex role of spatial context in variant pathogenicity is reflected from the difference in the classification rates of pathogenic residues in different regions. Consequently, this makes FunC-ESMs as a useful tool for providing mechanistic explanations of the effects of protein variants and their role in disease pathogenicity.

## Conclusions

Proteins in the human body interact with each other to fulfil their functions. A single amino acid substitution in a protein can drastically affect the way the protein interacts with other proteins, catalyses reactions, or binds substrates, and thereby directly alters function; other changes may compromise stability which in turn often leads to loss of function. Some of these changes can lead to disease, but the molecular mechanisms differ between different variants. Understanding how a protein loses its function provides useful information to understand how it works, how diseases occur, and how to potentially treat them. For example, identifying which variants cause loss of function via decreased stability and degradation is important for discovering the variants that might be treated using chemical chaperones or other perturbations of the cellular proteostasis machinery (***Bernier et al., 2004***; ***Nielsen et al., 2020***).

Previous experimental and computational studies have suggested that decreased protein stability is a common cause for how missense variants cause loss of function (***Wang and Moult, 2001***; ***Ferrer-Costa et al., 2002***; ***Yue et al., 2005b***; ***Casadio et al., 2011***; ***Berliner et al., 2014***; ***Gao et al., 2015***; ***Sahni et al., 2015***; ***Taipale, 2019***; ***Cagiada et al., 2021***, ***2023***). Therefore, we here set out to separate variants that cause loss of function via this mechanism version other ways that a missense variant may lead to perturbed function. To enable proteome-wide analysis, we leveraged accurate and computationally efficient sequence- and structure-based machine-learning methods to build FunC-ESMs. We validated FunC-ESMs and showed how it provides comparable or even improved accuracy to previous approaches (***Cagiada et al., 2023***). We applied FunC-ESMs to nearly all human proteins to shed light on the molecular mechanism behind loss of function for missense variants. Focusing on variants in folded regions, we find that approximately half of variants are predicted to cause loss of function, with slightly less than half of those caused by loss of stability. These numbers are comparable to previous analyses that used different experimental and computational methods for a smaller number of proteins or variants (***Wang and Moult, 2001***; ***Ferrer-Costa et al., 2002***; ***Yue et al., 2005b***; ***Casadio et al., 2011***; ***Berliner et al., 2014***; ***Gao et al., 2015***; ***Sahni et al., 2015***; ***Taipale, 2019***; ***Cagiada et al., 2021***, ***2023***) that support the robustness of our analysis.

Our analysis also provides predictions for functionally important residues in all human proteins, providing useful starting points for detailed experimental studies (***Cagiada et al., 2023***) or for supporting other prediction tasks such as for protein-protein or proteinligand interactions. We show that these functionally relevant residues tend to co-localize in groups; information that may be useful for example for discovering and characterising novel active sites (***Riziotis and Thornton, 2022***). This finding is consistent with previous work (***Jack et al., 2016***; ***Mayorov et al., 2019***; ***Hou et al., 2023***) and shows that many functionally relevant residues cluster around pockets, grooves or patches that enable them to perform their specific interaction. We also expect that our analysis of variant effects and molecular mechanisms will aid in interpreting experimental studies that assess variant effects at the proteome level (***Després et al., 2020***; ***Jun et al., 2020***; ***Hanna et al., 2021***; ***Cuella-Martin et al., 2021***).

Identifying whether a variant triggers loss of function, as well as identifying the underlying mechanisms, is important for determining if and how protein functionality could potentially be restored. Such knowledge can for example serve as a starting point for the development of therapies targeted at the specific mechanism. In our analysis of missense variants from ClinVar, we found that FunC-ESMs performed well in classifying variants, with benign and pathogenic variants classified correctly in most cases (91% and 79%, respectively). The percentage of pathogenic variants mislabelled as WT-like by our model might be attributed to several factors including (i) the higher rate of mispredictions for IDRs and (ii) that not all variants are loss of function (***Backwell and Marsh, 2022***) including for example variants that exert dominant-negative or gain-of-function effects that are difficult to predict using current computational prediction methods (***Gerasimavicius et al., 2022***). Finally, the balance between the fraction of true positives and false negatives is of course set by our specific choices of cut-offs, and could be tuned for individual proteins for specific purposes.

FunC-ESMs and the application to the human proteome offers a broad view of the underlying mechanisms of missense variants in the human proteome. We have here initially focused on separating a dominant mechanism (change in stability) from other mechanisms (perturbation of a broad class of functionally relevant residues). Future work would aim to expand this analysis into other more specific classes (***Arnaudi et al., 2022***). For example, with an increasingly accurate ability to predict structures of complexes with other proteins or nucleic acids at scale it should be possible to assess which variants perturb such interactions at the proteome level. Thus, with future advances in computational and experimental techniques our approach paves the way for more refined insights into the biology of human disease, potentially guiding the development of mechanism-based therapeutic strategies.

## Methods

### Sequence and structure of the target proteins

First, we first collected a list of UniProt IDs for the human proteome from the UniProt database (https://www.UniProt.org/help/api; organism = homo sapiens). We selected protein sequences present in the ‘one sequence per gene’ human reference proteome (UP000005640) from UniProt (2021_04) that was used for the AlphaFold Protein Structure Database (AFDB) (https://ftp.ebi.ac.uk/pub/databases/alphafold/v2/; ***Varadi et al. (2022b)***). Hence, proteins smaller than a minimum length of 16 amino acids, proteins containing non-standard amino acids, or proteins not in the UniProt reference proteome ‘one sequence per gene’ fasta file were excluded from our predictions. Structure predictions of the human proteome in the AFDB for proteins exceeding 2,700 amino acids are segmented into 1400 amino acids fragments, with a 1200 amino acid overlap between fragments; therefore we excluded these from our target protein list. Finally, as our variant effect predictions with ESM-1b were run against the UniProt 2023_02 version, all mismatched proteins between the two UniProt versions were removed. As a result of this filtering, our human proteome protein data set contained 20,144 unique unfragmented (i.e. shorter than 2700 amino acids) proteins.

### Sequence-based variant effect scores using ESM-1b

We used ESM-1b (***Rives et al., 2021***) to score the effect of variants on the target proteins. We used the pre-trained version of ESM-1b with 650 million parameters (ESM-1b_t33_650M_UR50S from https://github.com/facebookresearch/esm) with a masked marginal approach to evaluate the variant effect scores. Specifically, we mask the target position, evaluate the predicted probabilities for all amino acid tokens and calculate the score as the ratio between the probabilities of the wild-type amino acid and the target substitution. Because of memory and computational requirements for ESM-1b, there is a limit to the maximum length of sequence that can be analysed (1022 amino acid residues). To overcome this limit, we used a previously proposed strategy (***Brandes et al., 2023***) where proteins with more than 1022 amino acids were fragmented to fit into the system memory. We used a sliding window approach with 50% overlapping fragments to completely cover the sequence. The final scores were evaluated with a weighted average of the scores of the fragments. The weights are uniform for the position far from the termini and constructed with a sigmoidal function in window edge regions to mitigate potential edge effects.

### Structure-based variant effect scores using ESM-IF

We used ESM-IF (‘Inverse Folding’) to evaluate the structural effects of variants in a target protein. We used the pre-trained version of ESM-IF (esm_if1_gvp4_t16_142M_UR50 from https://github.com/facebookresearch/esm) with AF2 structures as input to generate and extract, from the ESM-IF language model decoder, amino acid conditional likelihood probabilities for each residue. We used these likelihoods to evaluate a variant score with the masked marginal approach as described above for ESM-1b.

Similar to the ESM-1b model, the ESM-IF language model can handle proteins up to 1022 residues. However, unlike ESM-1B, we noticed a flattening of the variant landscape score for proteins with more than 500 residues, leading often to incorrectly labelled variants near the protein C-terminus. We therefore used a fragmentation process (with a fragment of size 500) to process proteins that exceeded 500 residues. For each sequence fragment, we generated a new input structure consisting of: (i) a main chain containing all the residues of the selected fragment and (ii) an auxiliary chain containing the residues not present in the selected fragment. We used the structure as input for ESM-IF, using the main chain as the target for prediction and the auxiliary chain as a chain partner using the complex option. In this way, the ESM-IF graph can encode the complete information of the full structure and relay it to the language model, thereby producing high-quality predictions even for larger systems. We then used a sliding window approach to generate a final variant score by selecting the lowest score for the variant between the fragments in which the variant was present.

### Structural descriptors

We also calculated several structural descriptors for the proteins we analysed including different measures of exposure. We calculated the Accessible Surface area (ASA) and relative Accessible Surface Area (rASA) using DSSP from BioPython v.1.81 with default settings (***Rost and Sander, 1994***). We defined buried and exposed regions using an rASA cut-off of 0.20. We also calculated the half-sphere exposure (HSE, ***Hamelryck*** (***2005***)) for each residue to assess contacts in the direction of the side chain (‘up’) and contacts in the direction of the backbone (‘down’); the HSE calculations were performed using CA-CB vectors with a CB radius of 12Å, utilizing the Bio.PDB.HSExposure.HSExposureCB v.1.81 package. We also collected previously published (***Alderson et al., 2023***) predictions of disorder using SPOT (***Hanson et al., 2017***) to classify residues into folded regions and IDRs. To label cysteines forming disulphide bridges, we extracted their spatial coordinates from the AF2 structure using the Biotite python package and then applied two selection criteria for each pair of cysteines present: (i) the proximity of their two sulphur atoms within 2.05 ± 0.50 Å and a matching direction for their dihedral angles C*_β_* − S*_γ_* − S*_γ_* − C*_β_* within 90° ± 20°. When these criteria were met, we flagged both the cysteines as potentially forming a disulphide bond. In instances of multiple positive matches, we selected the pair with distance and dihedral angle closest to the optimal values, ensuring that each cysteine residue is predicted to form at most a single disulphide bond.

### FunC-ESMs

In FunC-ESMs we use a threshold-based model to classify variants in a target protein based on their mechanism of action. FunC-ESMs first separates ‘WT-like’ variants from deleterious variants based on their ESM-1b score. For those variants identified as deleterious, the ESM-IF score is then used to assign the ‘total-loss’ label to those that cause structural destabilisation or the ‘stable-but-inactive’ label to those that affect function but not stability. To this end, for both ESM-1b and ESM-IF, we needed to select a threshold to separate the different classes.

We used data generated by MAVEs included in the 2022 version of ProteinGym (***Notin et al., 2022***) to select the threshold for ESM-1b. Starting from the full dataset, we first removed MAVEs of ‘viral’ proteins and MAVEs with experimental protocols: ‘FACS’, ‘protein stability’ or ‘fluorescence’. To ensure maximal compatibility between the assay used in the MAVE and the prediction scores we selected only data from MAVEs that have Spearman’s correlation coefficient greater than 0.4 for ESM-1b and 0.3 for ESM-IF (Fig. S1A). This reduced the number of MAVEs from 87 to 32. Using the binarised scores present in ProteinGym, we performed a ROC analysis (python scikit-learn v1.0.2) using the ESM-1b data, and used the maximum of Youden’s index to select -6.5 as the threshold for ESM-1b (Fig. S1C).

We used ΔΔ*G* predictions on over two hundred thousand variants (***Tsuboyama et al., 2023***) to select a threshold for ESM-IF. From the full dataset, we excluded ‘de-novo’ proteins and further analysed only effects of single amino acid substitutions, resulting in a dataset of 173,684 ΔΔ*G* values. After confirming the correlation between the ESM-IF scores and the experimental ΔΔ*G* (Fig. S1B), we binarised the ΔΔ*G* predictions using -2 kcal/mol as a threshold, which has been shown in previous work to be a useful estimate to separate unstable from stable variants (***Jepsen et al., 2020***; ***Cagiada et al., 2021***; ***Høie et al., 2022***)). We then used a ROC analysis of the ESM-IF scores and the binary classification of the experimental data to evaluate the maximum of Youden’s index and -7 as the threshold for ESM-IF (Fig. S1D).

We used FunC-ESMs variant classification to assign a class to each residue in the human proteome. To this end, we labelled a residue as ‘tolerant’ if more than half of the variants at that position were WT-like, ‘structurally critical’ if more than half were totalloss, ‘functionally relevant’ if more than half were stable-but-inactive, or ‘mixed’ if none of the variant classes represent at least half of the substitutions at the target position.

### Radial distribution functions

We calculated the radial distribution function *g*(*r*), for different subsets of residues across all proteins. The radial distribution function *g*(*r*) for a given distance *r* is defined as:

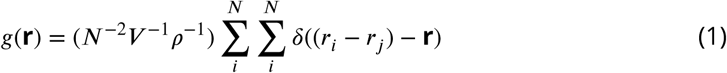

where *N* is the number of particles involved, *-* is the volume of the spherical shell and *-* is the density of the system. To obtain *g*(*r*) profile for a specific group of residues in a protein, we first calculated the distance between all the centres of mass of all the residues in the group using MDanalysis 2.5.0 (***Michaud-Agrawal et al., 2011***). We then selected a residue and binned the distances using spatial shells of thickness *r* + *dr*, where *dr* = 0.1 Å, around the target residue. We then normalised the counts in each bin (i) by the total number of particles (*N*^2^), (ii) by the volume (*-* = 2*nr*^2^*dr*) of the shell considered, and (iii) by the density of the system, evaluated as the ratio between the total number of positions considered and the volume of a sphere with the radius of gyration of the protein.

We analysed the result by separating the proteins based on their sequence length and for each group we evaluated a single representative *g*(*r*) by calculating for each bin the mean *g*(*r*) of all proteins present in the subgroup.

### Clinical annotations

We extracted the missense variants from ClinVar (***Landrum et al., 2014***, ***2017***) that have been categorized as (likely) benign, (likely) pathogenic, or variants of uncertain significance (VUS), the latter indicating that the pathophysiological consequences of the variant are not clear (***Landrum et al., 2017***). We collected this data in an in-house database according to previous protocols (***Tiemann et al., 2023***) by parsing the file from https://ftp.ncbi.nlm.nih.gov/pub/clinvar/tab_delimited/variant_summary.txt.gz (July 2023), and restricting the entries to those with a rating of at least one star, single-nucleotide variants, and were mapped to GRCh38 (script accessible at https://github.com/KULL-Centre/PRISM/tree/main/software/make_prism_files, release-tag v0.1.1) All ‘likely benign’ and ‘likely pathogenic’ variants were for this paper analysed together with ‘benign’ and ‘pathogenic’ variants, respectively. Variants with categorical labels other than these classes were discarded in the analysis along with all synonymous variants. To ensure that each variant was only represented once, we removed duplicate variants that had different ClinVar IDs but shared the same review status and clinical significance, retaining only a single entry for each unique variant. In this way we obtained 102,056 clinically annotated variants (69,909 benign and 32,147 pathogenic) in 12,114 unique human proteins.

## Data availability

The input data for the model and the predictions used in this study are available at the Electronic Research Data Archive at University of Copenhagen (KU/UCPH) (ERDA) and available via https://sid.erda.dk/cgi-sid/ls.py?share_id=DUWFpyjZp0. Our results can also be accessed via a Google Colaboratory (Colab) notebook available via https://github.com/KULL-Centre/_2024_cagiada-jonsson-func. UniProt ID codes for the proteins studied are available in a Text File format at this link.

## Code availability

The code used to generate the main text figures, to recreate the predictions, and generate new predictions is available via https://github.com/KULL-Centre/_2024_cagiada-jonsson-func. The repository also includes an implementation of the model that can be run using Google Colaboratory. We also provide a Colab notebook to assist accessing and downloading individual files from the data repository linked above, and for analysing and plotting data, helping with usability of our resources.

## Acknowledgments

We thank Kristoffer E. Johansson for assistance in collecting the SPOT-Disorder predictions. We thank Amelie Stein and Rasmus Hartmann-Petersen for many valuable discussions and input to our work. We thank the BioIcons (https://bioicons.com/) community for providing high quality icons for our figures and especially to Noelia Ferruz for the machine-learning icons used in Figure 1, which are licensed under CC-BY 4.0 Unported. We thank Andrew White for the protein emoji used in Figure 1 (licensed under CC-BY 4.0). The research was supported by the PRISM (Protein Interactions and Stability in Medicine and Genomics) centre funded by the Novo Nordisk Foundation (NNF18OC0033950, to K.L.-L.). We acknowledge access to computational resources via a grant from the Carlsberg Foundation (CF21-0392), and from the Biocomputing Core Facility at the Department of Biology, University of Copenhagen.

## Author contributions

Conception and design: MC, NJ and KL-L. Collection of data: MC and NJ. Analysis and interpretation of data: MC, NJ, and KL-L. Reformatting and data availability: NJ. Supervision: KL-L. Manuscript preparation: MC, NJ, and KL-L. Both MC and NJ contributed equally and have the right to list their name first in their CV.

## Supplementary Material

### Supplementary Figures

**Figure S1.**
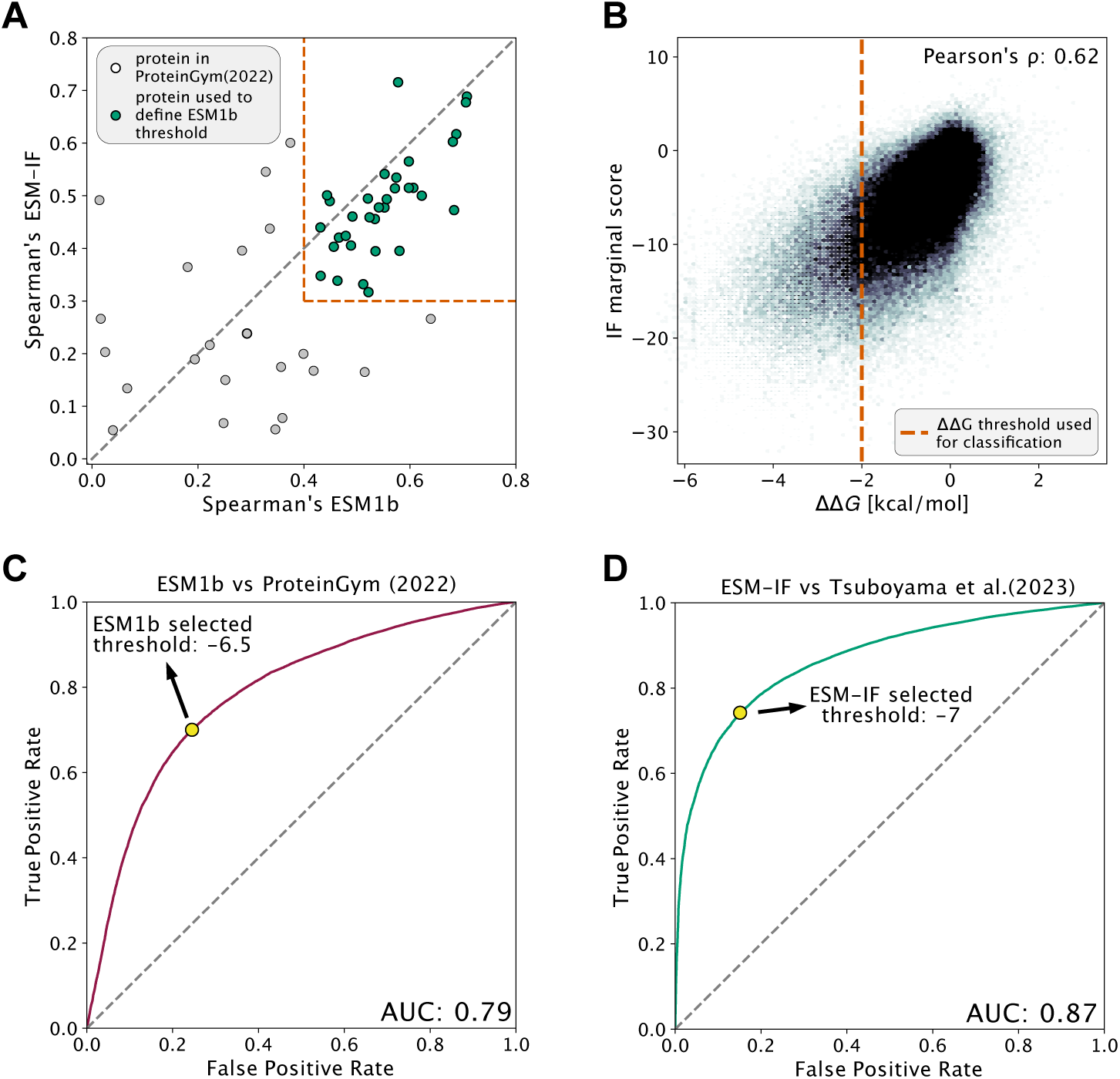
Details of the threshold selection for the FunC-ESMs model. (A) Scatter plot with Spearman’s correlations between the predictions of ESM-1b (x-axis) and ESM-IF (y-axis) against the experimental result for proteins included in the ProteinGym database. The red dotted lines indicate the systems used (in green) to select the threshold for ESM-1b. (B) Comparison between ESM-IF variant scores and ΔΔ*G* measurements of over 200,000 variants (***Tsuboyama et al., 2023***). The red dotted line represents the threshold that we used to define destabilized variants. (C and D) Area under the ROC curve used to select the thresholds for the two ESM predictors used in our FunC-ESMs model. In each plot, the AUC is shown in the bottom right corner and a yellow dot indicates the true positive and false positive coordinates for the threshold we selected by the maximal Youden’s index.

**Figure S2.**
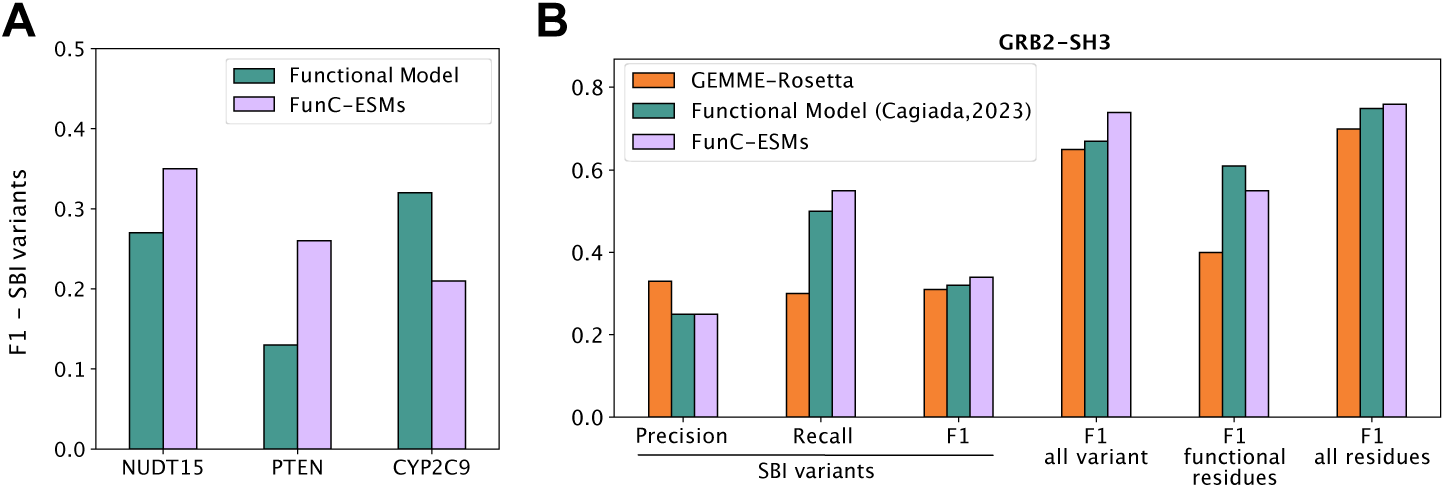
Validation FunC-ESMs and comparison to other methods. (A) Comparison for predictions of stable-but-inactive (SBI) variants between FunC-ESMs (lavender) and the predictions of the Functional Model (***Cagiada et al., 2023***) (green) on the proteins part of the training dataset used to train the Functional Model. To make the comparison fair, we retrained the Functional Model without the selected protein. (B) Comparison between FunC-ESMs (lavender), and a classification based on GEMME and Rosetta (***Cagiada et al., 2023***) (orange) and the Functional Model (***Cagiada et al., 2023***) (green) in predicting the variant mechanism and residue classes. The dataset used to compare the models is from an experimental classification on the GRB2-SH3 system. The y-axis shows the value for each metric used (and defined on the x-axis for each bar series).

**Figure S3.**
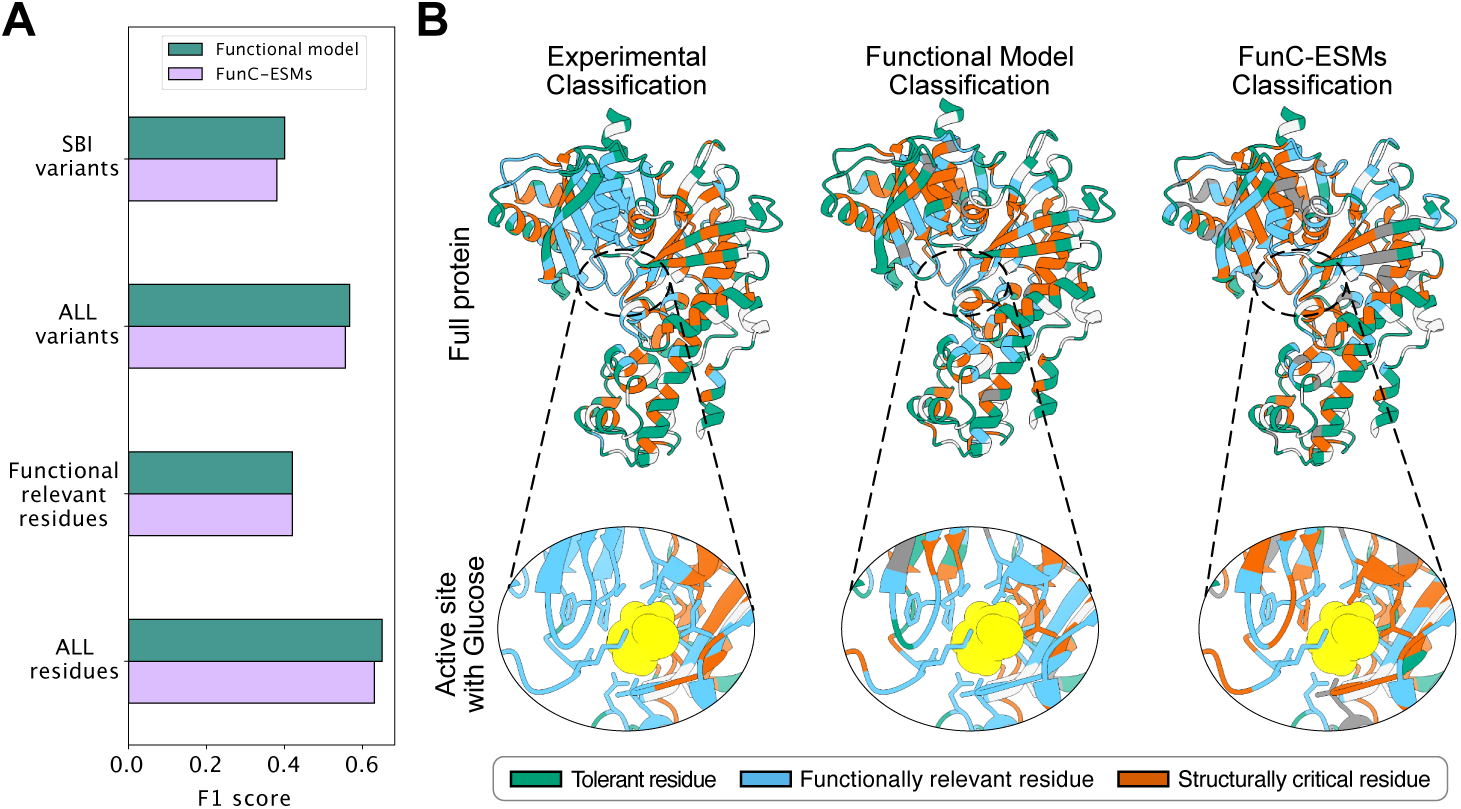
Validation of FunC-ESMs on data on glucokinase. (A) Comparison between predictions from FunC-ESMs (lavender) and the Functional Model (***Cagiada et al., 2023***) (green) and experimental data human glucokinase (***Gersing et al., 2023a***,b). (B) Cartoon visualisation of the classification using the experimental data (left), the Functional Model (centre) and FunC-ESMs (right) for human glucokinase. The upper part of the panel shows the full structure of glucokinase, with residues coloured according to the resulting residue class; a white colour indicates residues that were not classified in the experiment. The lower panel shows the area around the active site, including the glucose molecule (in yellow) and all side chains of residues closer than 7 Å.

**Figure S4.**
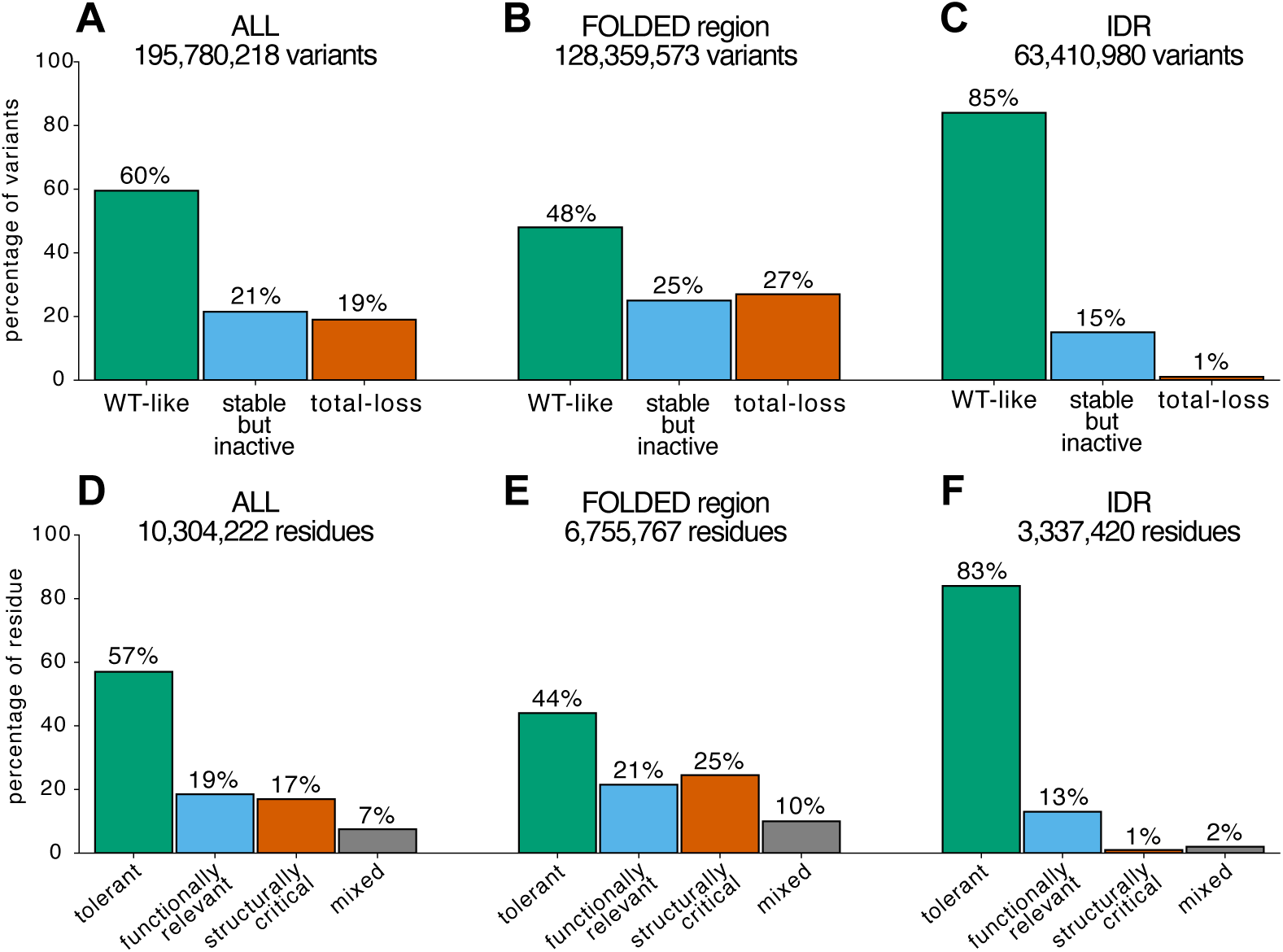
Percentages of variants and residues in each of FunC-ESMs’ class. Distribution of (A, B, C) variants and (D, E, F) residues for (A, D) all residues, (B, E) residues in folded regions and (C, F) residues intrinsically disordered regions of the human proteome.

**Figure S5.**
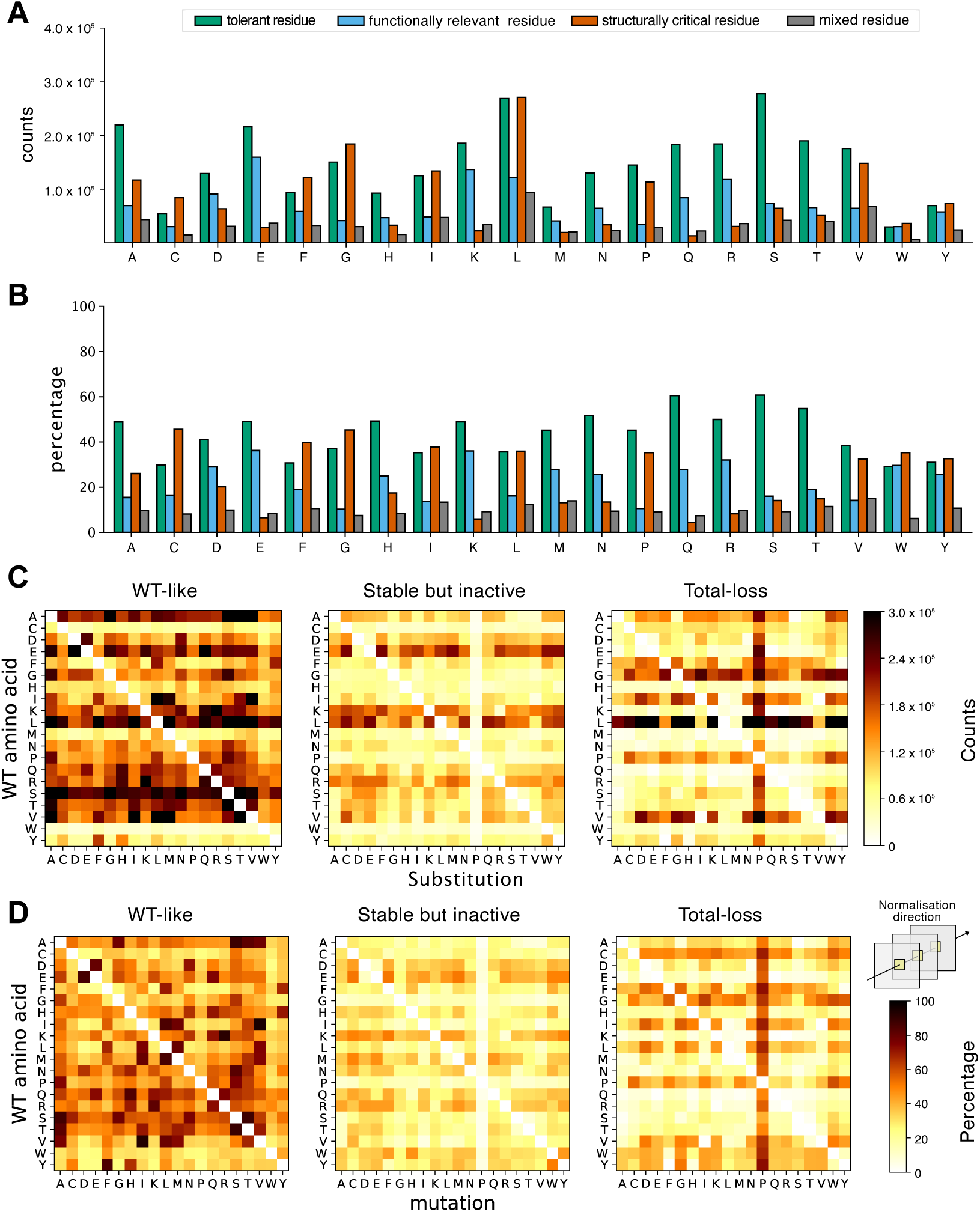
Contribution of different amino acid types and substitutions to the different predicted classes in folded regions. For each amino acid type, we present (A) the overall and (B) the normalized (by residue frequencies in the folded regions) counts for how each amino acid type contribute to the four position classes in the FunC-ESMs predictions. For each type of substitution (pair of ‘start’ and ‘end’ amino acid type), we show (C) the raw counts and (D) the normalized counts of variants falling into the three possible variant classes. The counts in (D) are normalised to the ‘start/end’ amino acid pair across the three different output labels.

**Figure S6.**
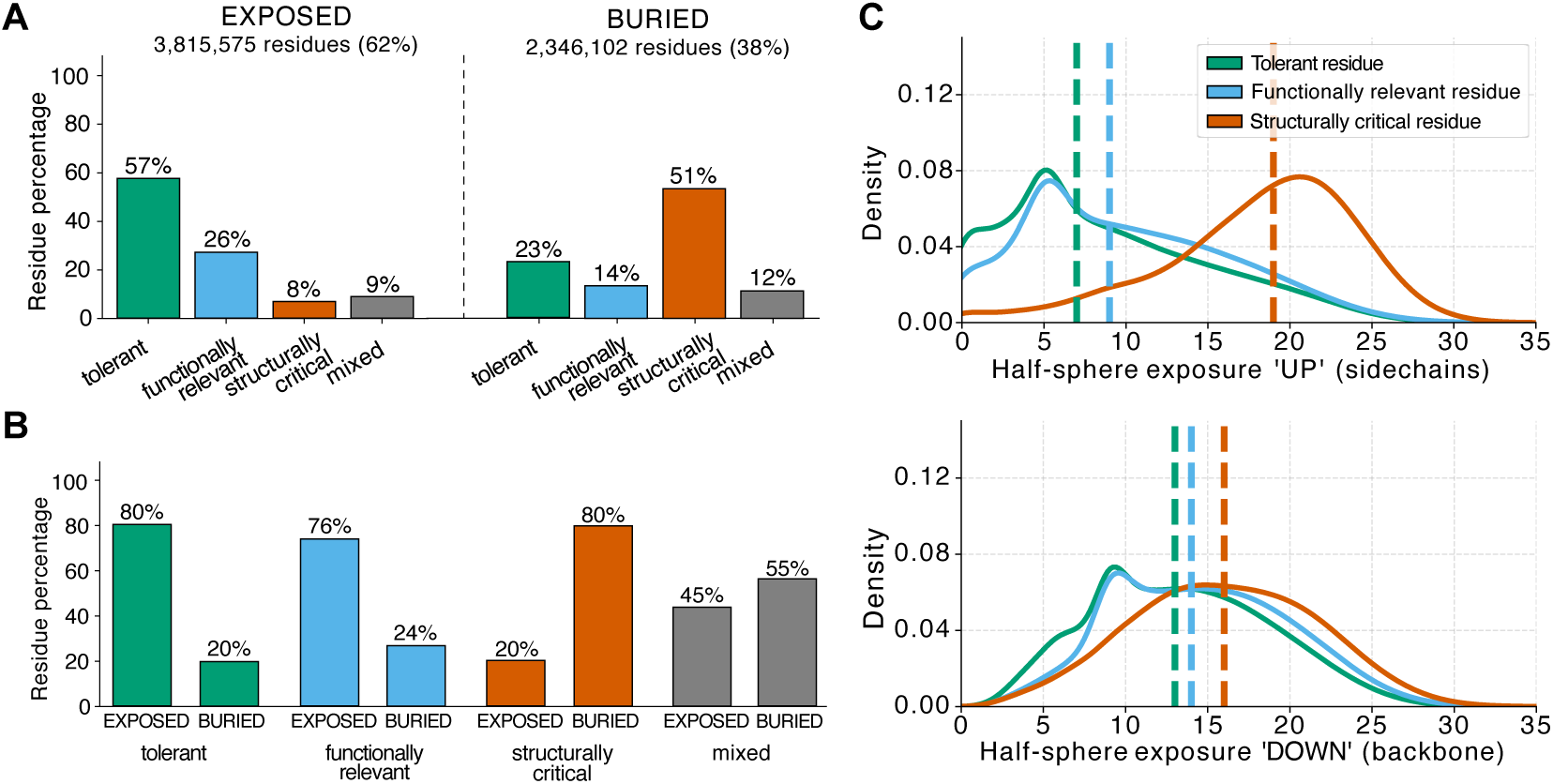
Statistics on solvent exposure and secondary structure elements in the folded domains of the human proteome. (A) Percentages of residues for each of the FunC-ESMs classes in exposed and buried regions and (B) percentages of each of the four residue classes in buried or exposed region. (C) Cumulative distribution of the half-sphere exposure components, with the up-component representing the contacts mostly surrounding the side chains and the down-component representing the direction of the backbone. The median value for each class is reported as a dashed line.

## Notes

https://github.com/KULL-Centre/_2024_cagiada-jonsson-func/

https://sid.erda.dk/cgi-sid/ls.py?share_id=DUWFpyjZp0

### Competing Interest Statement

KL-L holds stock options in and is a consultant for Peptone Ltd.

### Summary of Updates

Updates to how ESM-IF scores are calculated and to selection of variants from clinvar

